# Structural and Molecular Dynamics Analysis of Human β-Defensin 1 Variants Reveals Distinct Perturbation Signatures

**DOI:** 10.64898/2026.10.07.757448

**Authors:** Rebekah J. Fogarty, Raul Urrutia, Filippo Pinto e Vairo, Johnny C. Hong, Gwen Lomberk

## Abstract

Human β-defensin 1 (hBD-1) is a small, disulfide-constrained antimicrobial peptide produced predominantly in the kidney and urogenital tract, where it contributes to epithelial innate defense. Genetic variation in the gene encoding hBD-1 has been associated with altered expression levels and susceptibility to infections, including infection following solid organ transplantation, yet the molecular consequences of coding substitutions in the active, mature form remain largely undefined. Here, we integrate conventional genomics, protein bioinformatics, structural modeling, and molecular dynamics to investigate the molecular consequences of 93 human-observed hBD-1 missense variants. Evaluation of hBD-1 missense variants across sequence-, structure-, and dynamics-based dimensions reveals variant-specific molecular perturbation signatures that complement conventional and static biophysical predictions. By treating naturally occurring substitutions as a perturbation series across a compact defensin scaffold, this study identifies multiple molecular routes through which sequence variation can remodel an endogenous antimicrobial peptide and prioritizes variants for direct structure-function testing. These findings provide a mechanistic framework for investigating how naturally occurring hBD-1 variation may influence peptide behavior and, ultimately, antimicrobial function.

## Introduction

The kidney and urinary tract are normally protected, in part, through the production and secretion of antimicrobial peptides that contribute to epithelial host defense [1,2]. Disruption of this defense system may compromise urinary tract homeostasis and increase susceptibility to infection or disease. Genetic variation in innate immune effector pathways may modify host-defense capacity and contribute incrementally to susceptibility within complex clinical environments [3,4]. This may be particularly relevant after transplantation when host-defense capacity may be reduced by pharmacological immunosuppression, allograft-specific factors, and environmental exposure [4,5].

Defensins are a family of small, cationic, cysteine-rich antimicrobial peptides that contribute to this first line of innate host defense. Each has a predominantly β-sheet structure stabilized by three intramolecular disulfide bonds, and the disulfide-bridging pattern divides the family into α- and β-defensins [6,7]. More than 28 defensin genes have been identified in humans, with expression in the kidney, skin, cornea, and mucosal epithelia of the digestive, respiratory, and reproductive tracts [1]. Human β-defensin 1 (hBD-1) and human β-defensin 2 (hBD-2) are among the best characterized but differ substantially in their regulation. hBD-2 was first isolated from psoriatic skin and is induced by microbial or inflammatory signals, whereas hBD-1 is constitutively expressed and provides a more sustained component of epithelial antimicrobial defense [1,6–8]. hBD-1 was first isolated from hemofiltrate of patients undergoing chronic hemodialysis, supporting the kidney as a major site of expression [9].

hBD-1 is encoded by *DEFB1* and localizes to the loops of Henle, distal tubules, and collecting ducts, with the peptide also detectable in human urine [1,2]. Urinary hBD-1 concentrations have been found to increase during pyelonephritis [10], while reduced urinary and renal tissue hBD-1 abundance has been reported during acute kidney rejection after transplantation [11]. This small protein is synthesized as a 68-amino-acid precursor and undergoes N-terminal processing to generate a mature peptide, whose compact defensin fold is stabilized by three characteristic disulfide bonds [1,10,12,13] (Figure 1A). Importantly, hBD-1 antimicrobial activity is sensitive to its molecular state, with reduction of these disulfide bonds markedly altering antimicrobial properties [14]. Once secreted, hBD-1 functions through complementary antimicrobial and immunomodulatory mechanisms, including direct antimicrobial activity and chemokine receptor-mediated pathways [12,15]. Site-directed mutagenesis has identified residues in the cationic C-terminal region as important determinants of antimicrobial activity [12]. Together, these antimicrobial and immunomodulatory activities establish hBD-1 as an important component of innate host defense.

**Figure 1.**
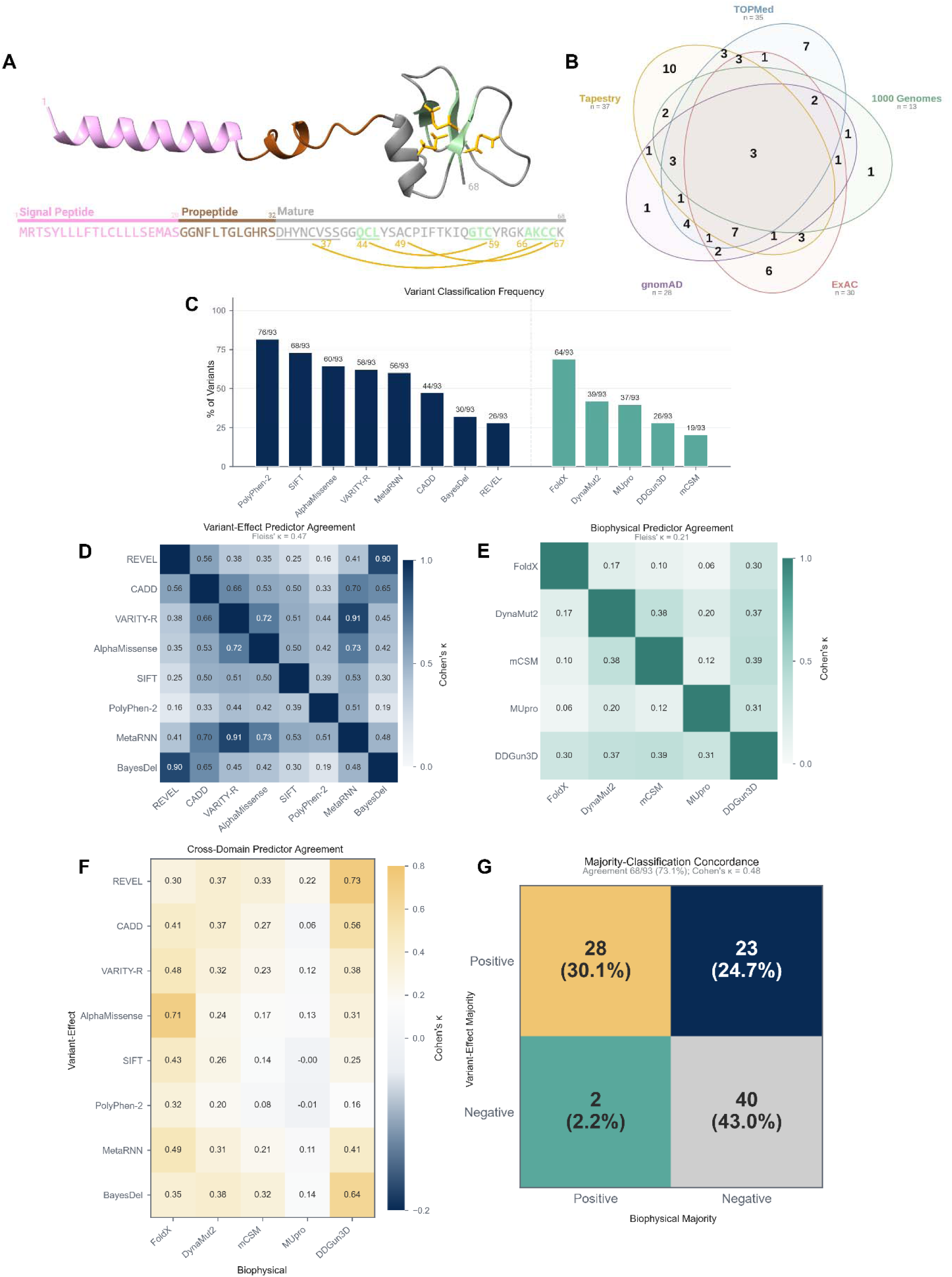
Variant landscape of human β-defensin 1 (hBD-1) and concordance across variant-effect and biophysical prediction domains. **(A)** Schematic representation of the hBD-1 precursor and mature peptide, showing the signal peptide, propeptide, mature region, and disulfide-bond connectivity. **(B)** Overlap of hBD-1 missense variants across major source datasets; numbers indicate variants within each intersection. **(C)** Frequency of positive classifications among the 93 analyzed variants for eight variant-effect predictors (blue) and five biophysical predictors (teal). Values above bars indicate the number of positive calls out of 93 variants. For biophysical predictors, a positive call denotes a predicted perturbation outside the predefined neutral range. **(D)** Pairwise Cohen’s κ agreement among variant-effect predictors. **(E)** Pairwise Cohen’s κ agreement among biophysical predictors shows lower overall concordance. Diagonal self-comparisons are not annotated in D and E. **(F)** Cross-domain pairwise Cohen’s κ values comparing each predictor. **(G)** Concordance between majority classifications from the two domains.

Non-coding *DEFB1* variation has been linked to infectious susceptibility in several settings, including severe sepsis [16] and invasive mold infections following solid-organ transplantation [17]. However, a direct relationship between mature hBD-1 coding variation and urinary tract infection or transplant outcomes has not been established [18]. This knowledge gap has become increasingly relevant as next-generation sequencing identifies numerous human coding variants whose molecular and functional consequences cannot be inferred directly from their occurrence in population or clinical sequencing resources.

Conventional missense-variant predictors provide an important first layer of annotation by integrating features such as sequence conservation, amino-acid substitution properties, structural context, and other genomic information. Structure-based biophysical stability predictors add information about the energetic consequences of amino acid substitutions within a defined structural context. Neither approach, however, directly captures the full range of time-dependent molecular behaviors that may be altered in a small, disulfide-constrained antimicrobial peptide. For hBD-1, these behaviors include flexibility, conformational organization, solvent exposure, secondary-structure maintenance, and intramolecular interaction networks. Because hBD-1 activity is closely linked to disulfide architecture, electrostatic properties, and conformational state, these additional features may be important for distinguishing how individual variants perturb peptide behavior.

We therefore assembled a panel of 93 human-observed single substitutions within the mature hBD-1 peptide and evaluated their molecular effects using an integrative approach combining conventional genomics, protein bioinformatics, structural modeling, and molecular dynamics simulations. We hypothesized that conventional variant-effect predictions provide an important foundation for variant characterization, while structural and dynamic analyses can reveal additional features relevant to hBD-1 peptide behavior, including perturbations in disulfide architecture, stability, flexibility, and intramolecular contacts. Rather than treating these approaches as competing measures of pathogenicity, we used them to interrogate distinct but complementary dimensions of variant effect. Our goal was to define variant-specific molecular perturbation signatures, identify variants whose dynamic behavior was not readily captured by static prediction approaches, and generate mechanistic hypotheses for subsequent functional investigation. Together, this approach was designed to provide a more resolved view of how coding variation may alter hBD-1 molecular behavior in biologically relevant renal and urogenital contexts.

## Materials and Methods

### Variant collection and curation

Mature hBD-1 missense variants were collected using UniProtKB/EBI entry P60022 for human β-defensin 1, encoded by *DEFB1* [19,20]. Variant annotations were obtained from the UniProt/EBI protein-variation record and included UniProtKB-curated variation as well as variants mapped from external genomic resources, including population reference panels, large-scale allele-frequency datasets, clinical variant databases, and registry resources. Sources included, among others, 1000 Genomes [21], ALFA and dbSNP [22], ClinVar [23], ESP [24], ExAC [25], Korea1K [26], Korea4K [27], ToMMo [28], TOPMed [29], and gnomAD [30]. Variants were mapped to the canonical hBD-1 protein sequence and restricted to missense substitutions within the mature hBD-1 peptide (residues 33-68). Synonymous, insertion/deletion, truncating, and somatic-only variants were excluded, along with variants that were mapped outside of the mature region, ensuring that the final panel comprised single amino acid substitutions in the mature peptide. Several variants carried somatic cancer-source annotations but were retained because they also had additional non-somatic source support.

The UniProt/EBI-derived variant set was supplemented with Mayo Clinic’s Tapestry cohort, a population genomics research cohort comprising approximately 97,000-100,000 individuals [31]. Thirty-seven mature hBD-1 missense variants in the final panel were observed in Tapestry, including three Tapestry-observed variants that were not present in the UniProt/EBI dataset at the time of curation. This source-driven curation strategy yielded a final primary panel of 93 mature hBD-1 missense substitutions for downstream analyses (Figure 1B). The panel was intended to capture human-observed mature-region hBD-1 missense variation across heterogeneous public annotation resources and Mayo Clinic patient cohort observations, rather than a cohort selected specifically for urinary tract or kidney disease.

### Computational variant-effect and biophysical impact predictions

Each missense variant was evaluated in the context of the canonical hBD-1 protein sequence (UniProt P60022). Eight computational variant-effect prediction algorithms were applied through the Ensembl Variant Effect Predictor (VEP, version 116) [32]: CADD [33], REVEL [34], SIFT [35], PolyPhen-2 [36], MetaRNN [37], BayesDel [38], VARITY [39], and AlphaMissense [40]. Because these algorithms differ in their underlying models and output formats, scores were interpreted according to published criteria for each method. Where necessary, continuous scores were converted to deleterious/pathogenic-likelihood categories using tool-specific published thresholds (Table 1). These classifications were treated as computational evidence of predicted variant effects, not as independent clinical determinations of pathogenicity.

**Table 1.** Predictor thresholds for binary classification.

| Domain | Predictor | Criterion counted as positive |
| --- | --- | --- |
| Conventional | CADD | $\geq 20$ |
| | REVEL | $\geq 0.50$ |
|  | SIFT | D (deleterious) |
|  | PolyPhen-2 | P or D (possibly/probably damaging) |
|  | MetaRNN | D (damaging) |
|  | BayesDel | D (damaging) |
| | VARITY-R | $\geq 0.50$ |
| | AlphaMissense | $> 0.564$ |
| Biophysical | FoldX | $< -0.5$ or $> +0.5$ |
| | DynaMut2 | $< -0.5$ or $> +0.5$ |
| | mCSM | $< -1.0$ or $> +1.0$ |
| | MUpro | $< -1.0$ or $> +1.0$ |
| | DDGun3D | $< -1.0$ or $> +1.0$ |

*Note: Biophysical predictors were evaluated using bilateral thresholds; both stabilizing and destabilizing changes exceeding the stated magnitude were counted as perturbing. Strict-majority classification required $\geq 5/8$ conventional predictors or $\geq 3/5$ biophysical predictors to be positive.*

Five computational approaches were used to estimate the biophysical effects of individual amino acid substitutions on structural stability using the resolved mature hBD-1 structure (PDB 1IJV) [41], expressed as changes in folding free energy (ΔΔG; kcal/mol): FoldX v4 [42], DynaMut2 [43], mCSM [44], MUpro [45], and DDGun3D [46]. Outputs were interpreted according to the published conventions and thresholds for each algorithm (Table 1). Because substantial deviation in either direction may reflect perturbation of the native energetic state, variants predicted to produce marked destabilization or stabilization were classified as having a predicted biophysical effect.

### Preparation of structures

The deposited AlphaFold full-length structure of human hBD-1 (AF-P60022-F1) was used directly as the structural starting model [47]. Residues 1-23 were removed, yielding a study construct comprised of residues 24-68 (Figure 1A). Although all variants examined in this study are located within the mature hBD-1 region spanning residues 33-68, the additional nine N-terminal residues (24-32) were retained to provide structural buffer between the peptide terminus and the variant-containing mature domain, thereby reducing the potential influence of terminal or truncation-related effects during molecular dynamics simulations. The experimental hBD-1 structure (PDB entry 1IJV) [41], used for biophysical structural impact predictions demonstrated high structural agreement with the AlphaFold model across the shared mature region (Cα RMSD = 0.55 Å), supporting consistency between the structural models used across analyses. Each missense substitution was introduced individually using MODELER implemented in Discovery Studio version 2026 [48,49]. Each mutated structure underwent side-chain rotamer refinement to resolve unfavorable steric contacts and optimize local starting geometry [50]. The three native hBD-1 disulfide bonds (C37-C66; C44-C59; C49-C67) were explicitly retained when unaltered. For cysteine-substitutions, the affected disulfide linkage was removed while the remaining native disulfide bonds were maintained. Molecular visualizations were generated using ChimeraX version 1.11 [51].

### Molecular dynamics (MD) simulations

#### Simulation preparation

The above-described variant structures underwent molecular dynamics (MD) simulations using Discovery Studio 2026 [49]. Ten independent simulations were performed for each variant and for wild-type (WT) hBD-1 using the CHARMm all-atom force field [52] and a distance-dependent implicit-solvent environment at pH 7.4. Independent replicates were initiated with random seeds. Each system underwent 1,000 steps of steepest-descent minimization, followed by 2,000 steps of adopted-basis Newton-Raphson minimization to relieve unfavorable interactions in the starting structure. Systems were then heated to 300 K over 50 ps and equilibrated for 300 ps before a 10 ns (10,000 ps) production simulation under an NVT ensemble. Atomic coordinates and trajectory frames were recorded every 100 ps. Trajectory stability was evaluated using potential energy (Supplementary Material, Figure S1). The final 2.5 ns, corresponding to the final 25 recorded frames of each trajectory, were used for subsequent analyses. Dynamic behavior was characterized by calculating root-mean-square fluctuation (RMSF), radius of gyration (R_g_), solvent-accessible surface area (SASA), and intramolecular contacts from the equilibrated portion of each trajectory. RMSF, R_g_, and SASA were calculated using all protein atoms. The same protocol was applied uniformly to WT and all variants so that the analysis remained comparative across the peptide panel rather than attempting to reproduce the full extracellular, membrane-associated, or receptor-bound environment of hBD-1.

#### Residue flexibility

Per-residue RMSF was calculated across the full simulated hBD-1 construct (residues 24-68) for each replicate system. For each residue, the difference between the mean variant RMSF and mean WT RMSF (ΔRMSF) was standardized by the residue-specific sample standard deviation across the 10 WT replicates, yielding a WT-standardized ΔRMSF value. Signed WT-standardized ΔRMSF values were retained to indicate the direction of flexibility changes, whereas absolute values were used to quantify perturbation magnitude. The magnitude component (S_RMSF(WT-std)_) was calculated as the mean absolute WT-standardized ΔRMSF across all 45 residues and scaled from 0 to 1, with a mean deviation of two WT standard deviations corresponding to the maximal score of 1. To additionally capture the spatial breadth of flexibility perturbation, residues with an absolute WT-standardized ΔRMSF ≥ 2 were designated as flexibility hotspots. The fraction of analyzed residues meeting this criterion was used to calculate the breadth component (S_RMSF(breadth)_), which was scaled such that hotspot classification of 25% or more of residues corresponded to the maximal score of 1. The overall RMSF perturbation score (*S_RMSF_*) was calculated as the equal-weight mean of the magnitude and breadth components:

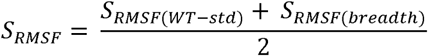

Further detailed equations are provided in Supplementary Material (S1-S8).

#### Conformational remodeling

Protein-level R_g_ and SASA were calculated across the converged simulation window using the MDTraj v1.11.1 Python package [53]. Protein Rg was calculated using MDTraj’s default implementation with an equal mass assigned to all protein atoms. SASA was calculated using the Shrake-Rupley algorithm [54] with a solvent probe radius of 1.4 Å and a sampling density of 960 surface points per atom. Values were averaged across the 10 independent replicates before comparison with WT. Mean differences were standardized by the corresponding WT standard deviations, and absolute standardized values were scaled from 0 to 1 using two WT standard deviations as the upper reference for perturbation (S_Rg_, S_SASA_). Because R_g_ and SASA capture complementary features of conformational remodeling, the two normalized measures were combined to generate an overall conformational-remodeling score:

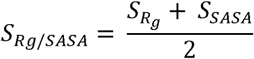

Further detailed equations are provided in Supplementary Material (S1-S3, S9).

#### Intramolecular contacts and native-fold integrity

Intramolecular noncovalent contacts, including hydrogen bonding, hydrophobic, and electrostatic interactions were quantified over the converged portion of each MD trajectory. Contact analysis was restricted to the variant position and its interacting residues to quantify remodeling of the local interaction network relative to WT. Contact occupancies were calculated for each variant residue-partner interaction and compared with those observed at the corresponding WT position. Differences in interaction profiles were summarized using a weighted Jaccard distance yielding a contact-remodeling score (S_contact_), ranging from 0, indicating identical interaction profiles, to 1, indicating complete divergence. Detailed equations are provided in Supplementary Material (S10-S11).

Native-fold integrity was evaluated independently by quantifying β-strand occupancy across each trajectory. The conserved antiparallel β-strand core has been implicated in defensin function [12,13]. Secondary structure was assigned for each WT and variant replicate using DSSP version 2.2.0 implemented through MDTraj [55,53]. For residues assigned to β-strands in the experimental structure (43-45, 57-59, and 64-67), β-strand occupancy was calculated across the analyzed trajectory window. Absolute differences in mean occupancy relative to WT were standardized by the WT standard deviation and scaled from 0 to 1 using two WT standard deviations as the upper perturbation reference, generating the secondary-structure perturbation score (S_ss_). Contact remodeling and secondary-structure perturbation were then combined to generate a fold-integrity perturbation score:

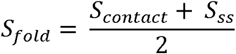

Further detailed equations are provided in Supplementary Material (S12-S14).

### Integration of variant-effect prediction, biophysical prediction, and MD analyses

Complementary scores were generated to summarize variant-associated effects within each analytical domain. For computational variant-effect prediction, a positive consensus call required a strict majority, defined as at least 5 of the 8 algorithms classifying the variant as deleterious or as having increased pathogenicity likelihood according to tool-specific thresholds. For biophysical prediction, a positive consensus call similarly required at least 3 of the 5 algorithms to identify the variant as producing a predicted biophysical perturbation according to the prespecified tool-specific criteria. These scores were retained as separate measures representing complementary dimensions of variant effect.

To quantify perturbations observed in the time-dependent MD simulations, an MD-derived dynamics score (*S_MD_*) was calculated as the equal-weight mean of three components: RMSF effect, R_g_/SASA remodeling, and intramolecular contact-network/fold alteration. Equal weighting was used as a descriptive integration strategy and was not trained or calibrated against antimicrobial activity or clinical phenotype:

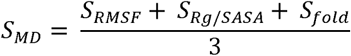

All component scores were normalized from 0 to 1, with higher values indicating greater modeled variant-associated perturbation within the corresponding analytical domain. Scores from the variant-effect prediction, biophysical prediction, and MD domains were retained separately to permit assessment of agreement or discordance among complementary measures of variant effect.

### Statistical analysis

Statistical analyses evaluated agreement among prediction methods, associations between computational domains, and differences in dynamic measurements. Predictor agreement was assessed using Fleiss’ kappa (κ) for comparisons involving more than two classifiers [56] and Cohen’s κ for pairwise comparisons [57]. Continuous domain scores were compared using Spearman rank correlation, and variant-versus-WT MD measurements were assessed using two-sided Welch’s t-tests. For MD measurements subjected to variant-wise hypothesis testing, p-values were adjusted separately within each metric using the Benjamini-Hochberg false-discovery-rate procedure, with q < 0.05 considered significant (Supplementary Material, Figure S4).

## Results

### Variant-effect predictors converge on conserved hBD-1 residues but differ across other variants

Eight computational variant-effect prediction algorithms were applied to the 93 hBD-1 variants to establish a reference profile of predicted effects (Figure 1C). Across the eight algorithms, moderate agreement was observed (Fleiss’ κ=0.473; Figure 1D). The tools produced unanimous classifications for 32/93 variants (34.4%), including 25/93 (26.9%) classified as deleterious or damaging by all algorithms. Using a strict-majority criterion (≥ 5/8 tools), 51/93 (54.8%) variants were classified as having a deleterious effect.

Unanimous deleterious classifications were observed for 19/20 cysteine substitutions and all (n=6) substitutions at G57. Notably, G57 was the only non-cysteine residue conserved across the β-defensin paralogs examined (Supplementary Material, Figure S2). In contrast, seven variants received unanimous benign predictions, representing substitutions involving serine, isoleucine, or arginine. Although not among the unanimously benign variants, V38I was classified as benign/tolerated by all variant-effect predictors except PolyPhen-2 and had a general-population allele frequency of 0.0075, with homozygous observations.

Cysteine substitutions showed particularly consistent predictions across algorithms. All 20 met the strict-majority damaging criterion, with C37W representing the only cysteine substitution not uniformly classified across all eight algorithms because of its CADD prediction. Despite the structural importance and strong conservation of disulfide-forming cysteine residues, several cysteine substitutions are present in population datasets. C67S, for example, was the second most frequent hBD-1 amino acid substitution in gnomAD and has also been observed in the homozygous state.

Individual predictors differed substantially in the frequency of positive classifications (Figure 1C). PolyPhen-2 was the most permissive, classifying 81.7% of variants as damaging, whereas REVEL was the most conservative, with only 28% of variants exceeding its pathogenicity-likelihood threshold. All REVEL-positive variants involved either a cysteine substitution or alteration of G57, indicating that REVEL preferentially identified substitutions at highly conserved positions within the hBD-1 panel. REVEL and CADD showed moderate pairwise agreement (Cohen’s κ = 0.56), and both showed low-to-moderate agreement with AlphaMissense (REVEL, κ=0.35; CADD, κ=0.53). These partial concordances show that established prediction methods can assign different effects to the same hBD-1 substitution because of differences in underlying features, model architectures, and decision thresholds.

### Biophysical predictors reveal heterogeneous stability effects across hBD-1 variants

Computational biophysical tools were used to evaluate predicted stability effects of the 93 hBD-1 variants relative to the WT structure. Agreement across the five algorithms was low (Fleiss’ κ=0.21), reflecting substantial heterogeneity in predicted biophysical effects (Figure 1E). This discordance occurred among both cysteine and non-cysteine substitutions. Using a strict-majority criterion (≥ 3/5 algorithms), 30/93 (32.3%) variants were classified as having a predicted stability perturbation.

Only 5/20 (25%) cysteine substitutions were predicted to perturb structural stability by all five algorithms. FoldX and DDGun3D classified all cysteine variants as biophysically perturbing, while mCSM identified substantial stability effects in fewer than half. FoldX produced the highest overall perturbation rate, classifying 68.8% of all variants as having altered stability, whereas each remaining algorithm classified fewer than 50% (Figure 1C). Agreement among individual biophysical predictors was generally low, although DDGun3D showed the highest overall concordance with the other algorithms. Fourteen variants received unanimous predictions of minimal or neutral stability effects. Overall, the biophysical predictors produced a heterogeneous landscape of predicted stability perturbation, distinct from the more concordant classifications generated by the variant-effect predictors.

### Concordance analysis identifies variants with discordant predicted effects

To compare classifications across the variant-effect and biophysical prediction domains, pairwise concordance was assessed across all 93 hBD-1 variants. A strict-majority criterion was applied within each domain to generate a binary classification for each variant. Cross-domain agreement varied substantially across predictor pairs, with the strongest concordance between AlphaMissense and FoldX (Cohen’s κ = 0.71) and between REVEL and DDGun3D (Cohen’s κ = 0.73). The two domains agreed for 68/93 variants (73.1%; Cohen’s κ = 0.480) (Figure 1F). Twenty-eight variants were classified as deleterious or biophysically perturbing by both domains, whereas 40 showed no predicted effect above predefined thresholds in either domain (Figure 1G). The remaining 25 variants showed discordant classifications; 23/25 (92.0%) were classified as deleterious by variant-effect predictors but did not meet the majority threshold for biophysical perturbation. These findings identify a subset of hBD-1 variants for which sequence-based and structure-based computational predictions provide distinct evidence regarding potential variant effects.

### hBD-1 variants exhibit local, proximal, and distal flexibility perturbations

Previous studies have shown that variant-associated changes in protein dynamics, including alterations in flexibility, global compactness, solvent exposure, and interaction networks, can provide mechanistic hypotheses of molecular perturbation [58–61]. Accordingly, we performed MD simulations for all 93 hBD-1 variants relative to WT to characterize changes in dynamic behavior. Initial analyses focused on residue-level flexibility, quantified by RMSF across replicate trajectories. Across the 93 variants, 74 (79.6%) exhibited a positive mean signed ΔRMSF relative to WT, indicating that the average flexibility change for most variants was toward increased flexibility (Supplementary Material, Figure S3A-B).

Sequence-distal substitutions can affect the flexibility and dynamic coupling of functionally important residues, providing a potential mechanism through which variants outside canonical functional sites alter characteristic peptide behavior [62]. To characterize the breadth of flexibility alterations caused by hBD-1 variants, the residues with absolute WT-standardized ΔRMSF differences ≥2 were classified as flexibility hotspots. Twenty-nine variants contained at least one hotspot, and several variants showed perturbations across multiple residues (Supplementary Material, Figure S3C). C37Y and G62A produced the broadest flexibility effects, with 13 and 9 hotspots, respectively (Figure 2A). All 9 hotspots identified from G62A were also present in C37Y.

**Figure 2.**
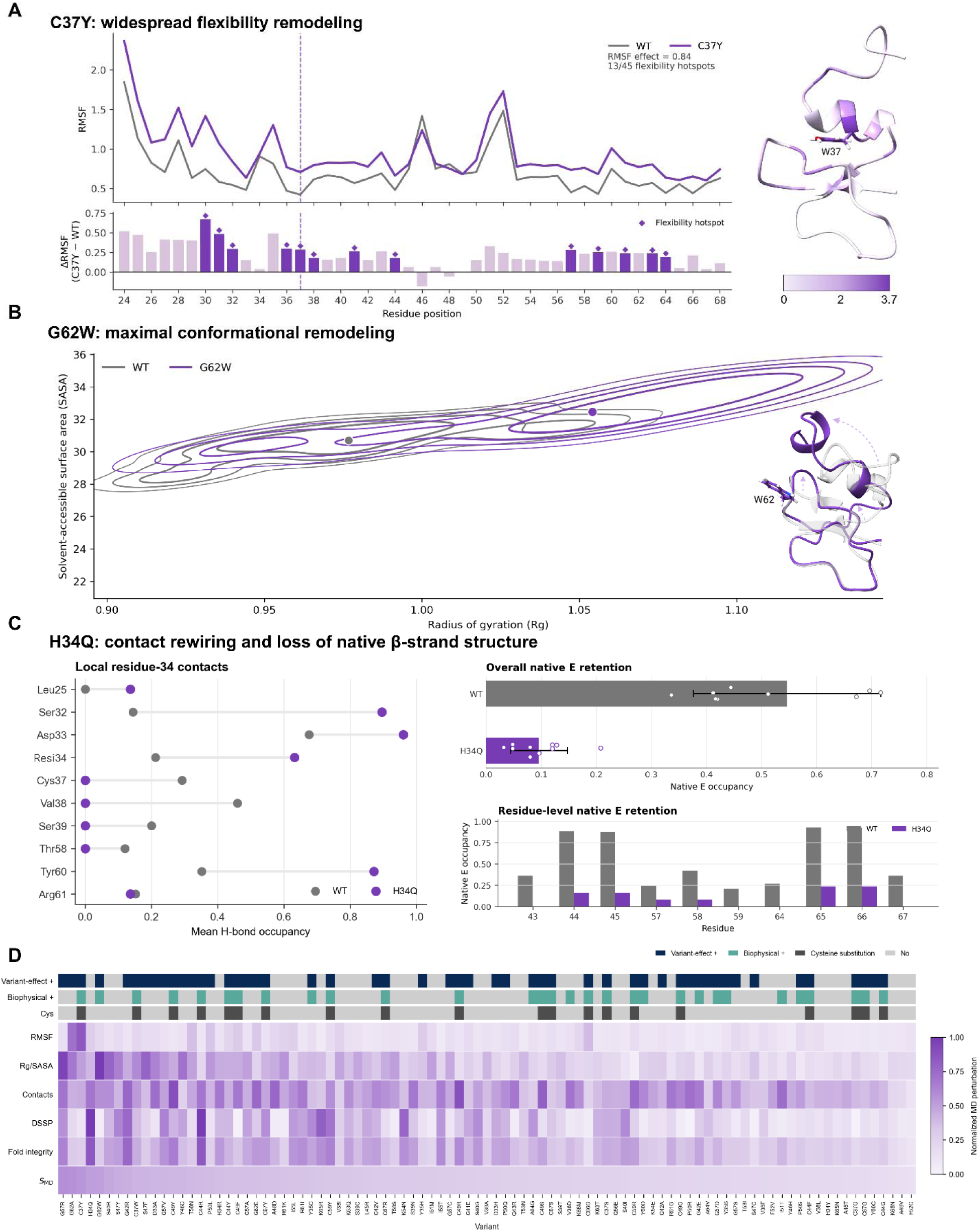
Molecular dynamics profiling reveals distinct mechanisms of variant perturbation. (A-C) Representative variants illustrate distinct modes of MD perturbation: **C37Y**, widespread flexibility remodeling, with the 3D structure depicting deviations from WT (WT-standardized |ΔRMSF|≥2 defining hotspots); **G62W**, maximal conformational remodeling in Rg/SASA space, with the 3D structure depicting regions of large deviation and expansion from WT; and **H34Q**, local contact rewiring with loss of native β-strand retention. **(D)** MD perturbation landscape across 93 hBD-1 variants ranked by overall *S_MD_.* The heatmap shows normalized RMSF, Rg/SASA, contact, DSSP, fold-integrity, and *S_MD_* scores; annotation tracks indicate domain-level variant-effect and biophysical classifications and cysteine substitutions.

Several variants, including S47F/Y, G62W, C37W, P50Q, and C44F, produced highly localized flexibility changes, with hotspots restricted to the substituted residue (Supplementary Material, Figure S3C-E). In contrast, recurrent hotspots were observed at positions distal to the substitution site, suggesting coordinated long-range changes in peptide dynamics. V38, for example, was found as a hotspot across 11 variants arising from eight distinct mutation sites, six of which involved substitutions located more than three residues away in the primary sequence (Supplementary Material, Figure S3D-E). These patterns are consistent with sequence-distal dynamic coupling within hBD-1.

To integrate the magnitude and spatial extent of flexibility perturbation, an RMSF-derived score incorporating mean flexibility perturbation and hotspot breadth (*S_RMSF_*) was calculated for each variant (Figure 2A,D). C37Y and G62A showed the largest effects, with scores approximately two-fold higher than the next-ranked variant. Most variants, however, produced limited large-scale flexibility changes, indicating preservation of the relatively rigid regions identified in WT hBD-1 (Figure 2D).

### hBD-1 variants differentially alter global compaction and surface exposure

To assess broader conformational effects, we quantified radius of gyration (Rg) and solvent-accessible surface area (SASA) across all 93 hBD-1 variants relative to WT. Rg evaluated overall molecular compaction, whereas SASA provided a complementary measure of surface exposure. Twenty-four variants showed nominally significant differences in Rg relative to WT (p<0.05), and nine showed nominally significant increases in SASA (Supplementary Material, Figure S4A-B). After Benjamini-Hochberg correction for multiple testing, Rg differences remained significant for S40R, G57R, and G62W. G62W and G57R showed the greatest conformational remodeling across the combined Rg and SASA measures (Figure 2B), whereas C59R showed the least remodeling across Rg and SASA, followed by P50S, G42E, C44G, and S40I. To summarize these complementary measures, a global conformational perturbation score (*S_Rg/SASA_*) was calculated for each variant (Figure 2D). Fifteen of 93 variants (16.1%) had scores greater than 0.5 on the normalized 0-1 scale. Several variants also showed discordant effects between Rg and SASA, indicating that changes in compaction were not uniformly accompanied by altered surface exposure. These patterns are consistent with variant-specific conformational rearrangements rather than a single mode of global structural disruption.

### Charge-altering substitutions show prominent local contact-network remodeling and β-core perturbation

To assess local structural effects beyond flexibility and global conformation, we characterized intramolecular interaction networks for each hBD-1 variant relative to WT. Both gains and losses of residue-level contacts were quantified to capture remodeling of the local interaction environment. Several positions showed marked contact-network remodeling across multiple substitutions (Supplementary Material, Figure S4C-D). Specifically, all five substitutions at C49 produced substantial local contact perturbation, consistent with the constrained structural environment surrounding this residue within the loop between the β1 and β2 strands. Similar effects were observed for C59 and C44 substitutions, while substitutions at the C-terminal cysteines C66 and C67 generally produced more modest contact rearrangements. Among non-cysteine substitutions, the largest contact-network perturbations were associated with changes in residue charge (i.e., introduction or loss of acidic/basic amino acids). Variants combining electrostatic and steric alterations, including A48D, G42R, and G62E, produced particularly strong remodeling of the local interaction network (Supplementary Material, Figure S4C-D).

Because the conserved antiparallel β-sheet is a central feature of the defensin fold, β-strand occupancy was evaluated throughout the MD trajectories at residues 43-45, 57-59, and 64-67. H34Q and C44R showed the strongest and most persistent reductions in native β-sheet occupancy across replicates, while 16 variants exhibited complete loss of native β-strand assignment for monitored residues in at least one replicate (Supplementary Material, Figure S4E). Thus, both cysteine and non-cysteine substitutions produced modeled losses of native β-strand occupancy, although the magnitude and consistency of these effects varied among variants.

To integrate local interaction remodeling with β-core maintenance, a fold-integrity perturbation score (*S_fold_*) was calculated as an equal-weight mean of the contact-network and secondary-structure components. H34Q and C44R exhibited the strongest overall fold-integrity perturbations, characterized by substantial changes in intramolecular contacts and β-sheet occupancy despite relatively modest effects on overall flexibility or global conformation (Figure 2C-D). More broadly, these analyses show that modeled structural perturbation can remain highly localized, with substantial changes in contact organization or secondary-structure maintenance occurring without correspondingly large changes in global compaction or solvent exposure.

### MD-derived perturbation scores provide mechanistic resolution across prediction profiles

An MD-derived dynamics score (*S_MD_*) was calculated to summarize variant-associated perturbations across flexibility, global conformation, and local secondary-structure/contact-network behavior (Figure 2D). Scores were normalized from 0 to 1, with higher values representing greater modeled perturbation. Variant-effect and biophysical scores were positively correlated (Spearman ρ = 0.68), whereas *S_MD_* showed little correlation with either the variant-effect (ρ = 0.07) or biophysical (ρ = -0.18) static domains (Figure 3A), indicating that *S_MD_* captures a largely distinct dimension of modeled variant perturbation. Variants with *S_MD_*at or above the 75th percentile of the analyzed distribution (Q3 ≈ 0.355) were designated high dynamics-perturbing variants for relative prioritization, identifying 24/93 variants (25.8%; Figure 3B). Among these 24 variants, 5 (20.8%) were classified as benign or neutral by both static prediction domains, indicating dynamic perturbations not reflected in either domain (Figure 3B). In addition, 11/24 (45.8%) high dynamics-perturbing variants were classified as deleterious by variant-effect predictors but did not meet the majority threshold for biophysical prediction algorithms. For example, G57R, which exhibited the highest *S_MD_* score, exceeded the positive threshold for all eight variant-effect predictors but did not meet the biophysical majority criterion.

**Figure 3.**
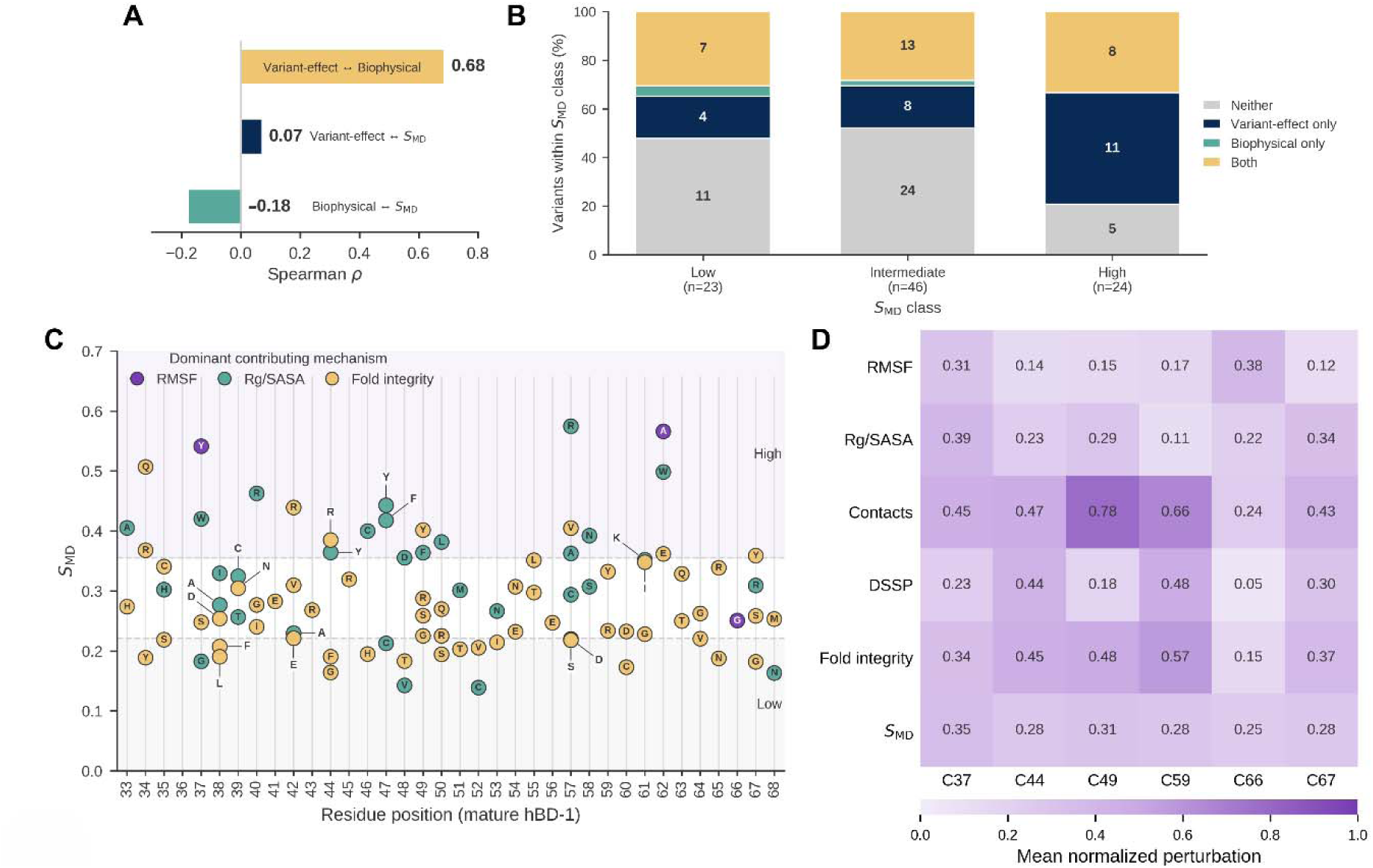
Integrated comparison of static prediction and MD perturbation across hBD-1 variants. **(A)** Spearman correlations among variant-effect, biophysical, and scores. **(B)** Distribution of static predictor support across low, intermediate, and high *S_MD_* classes. **(C)** Position-specific scores colored by dominant MD mechanism. **(D)** Mean normalized MD-component perturbation across native cysteine positions.

Variants with *S_MD_* values between the first- and third-quartile cutoffs (0.221≤*S_MD_*<0.355) were designated intermediate dynamics-perturbing variants, comprising 46/93 variants (49.5%). Of these, 24/46 (52.2%) were classified as benign or neutral by both prediction domains, while an additional 9 showed discordant classifications between the two domains (Figure 3B). Notably, incorporation of the dynamics-based analyses provided additional mechanistic insight into how these variants may perturb hBD-1 structure and function. Importantly, MD-derived perturbations were not limited to variants with discordant predictions. High or intermediate *S_MD_* scores also aligned with concordant deleterious or perturbing classifications from both the variant-effect and biophysical domains for an additional 21/93 variants (22.6%). Together, these findings illustrate how dynamics-based analysis can identify structural and conformational perturbations in variants that remain unresolved or appear neutral across static prediction domains.

Conversely, 23 variants fell within the low dynamics-perturbation group (*S_MD_*< 0.221). Of these, 16/23 (69.6%) lacked concordant damaging classifications by both static domains, including 11 variants (47.8%; A48T, A48V, F52C, F52V, H34Y, K65N, K68N, T53I, V38F, V38L, and Y46H) that did not satisfy the majority threshold in either domain (Figure 3B). This pattern demonstrates high cross-domain concordance among variants with limited modeled dynamic perturbation. Overall, *S_MD_* stratification captured a spectrum of variant-associated dynamic behavior, dependent on substitution position and replacement residue (Figure 3C), ranging from limited perturbation to substantial dynamic remodeling in variants with either concordant or discordant static classifications.

### Cysteine substitutions produce heterogeneous structural and dynamic perturbations

Cysteine substitutions showed substantial discordance across variant-effect, biophysical, and dynamics-based analyses. Given the central role of disulfide bonding in hBD-1 structure, we examined these variants in greater detail to determine whether disruption of individual cysteine residues produced uniform or distinct molecular effects. As previously described, variant-effect predictors showed near-complete agreement, with 19/20 cysteine substitutions unanimously classified as deleterious, and all 20 satisfying the strict-majority criterion. Although biophysical predictors showed greater heterogeneity at the individual algorithm level, all cysteine substitutions also met the majority criterion for predicted stability perturbation (Figure 2D).

In contrast, *S_MD_* stratification distributed cysteine variants across low (n = 4), intermediate (n = 9), and high (n = 7) dynamics-perturbation classes, with 16/20 (80%) showing at least intermediate perturbation (Figure 2D). Individual *S_MD_*components further demonstrated that cysteine variants affected distinct dynamic features, with relatively few causing simultaneous perturbation across all components. Thus, despite uniform majority support from the two static prediction domains, cysteine substitutions did not produce a uniform dynamic phenotype. Perturbation signatures varied according to both cysteine position and replacement residue (Figures 2D and 3D). Substitutions affecting the C37-C66 linkage were more strongly associated with flexibility changes, whereas substitutions involving the C44-C59 and C49-C67 linkages preferentially affected local secondary-structure maintenance and intramolecular contact organization (Figure 3D). Mean *S_MD_* also varied by replacement-residue class, with lower values for glycine or serine substitutions (0.221) than for arginine (0.304) or aromatic substitutions (0.372), suggesting that the biochemical properties of the substituted residue contribute to the magnitude of modeled perturbation within the compact β-core.

## Discussion

In this study, we define distinct molecular perturbation signatures across a broad set of human hBD-1 missense variants by integrating variant-effect prediction, structure-based biophysical analysis, and molecular dynamics simulations. These approaches interrogate complementary dimensions of hBD-1 peptide behavior and produce distinct patterns of predicted or modeled perturbation. This multidimensional analysis was particularly informative for variants with discordant prediction profiles, revealing changes in flexibility, conformational organization, intramolecular contacts, or secondary-structure integrity that were not apparent from sequence- or stability-based assessments alone. MD further provided time-dependent mechanistic resolution into how individual substitutions altered hBD-1 molecular behavior.

hBD-1 is an important component of innate defense in the kidney and urogenital tract, where preservation of peptide structure and molecular behavior is likely relevant to antimicrobial function. Human variation in *DEFB1* has been associated with infection-related phenotypes, including invasive mold infection following solid-organ transplantation [17], and altered hBD-1 expression has been reported in kidney transplantation. These observations support interpreting *DEFB1* variation within a host-susceptibility framework, in which individual alleles may modify antimicrobial capacity without independently causing disease. Modest molecular effects could become more consequential in settings of reduced host-defense capacity, including pharmacologic immunosuppression. Our analysis provides mechanistic insight into how individual amino acid substitutions may differentially perturb hBD-1 structure and dynamics. However, this study does not test clinical association and does not establish that any variant causes urinary tract infection, graft dysfunction, rejection, or other clinical outcomes. Instead, it defines molecular phenotypes that can be carried forward into functional assays and appropriately powered genotype-phenotype validation studies in renal, urogenital, and immunosuppressed settings.

An important implication of this study is that hBD-1 variant effects may not be described adequately by a single molecular property. Sequence conservation, energetic stability, local flexibility, conformational organization, and residue-level interaction networks represent distinct but interconnected features of peptide behavior. Discordance among these measures may therefore identify mechanistically different classes of variants. This is particularly relevant for a small, disulfide-constrained peptide such as hBD-1, in which a substitution may preserve overall compaction while altering local interactions or flexibility elsewhere in the structure. The individual domain scores should therefore be interpreted as complementary summaries of different molecular properties. Because predictors within a given domain share related features or assumptions, majority classifications reflect within-domain consensus and should not be considered fully independent lines of evidence.

MD simulations added a time-dependent view of these effects under the modeled conditions. Their value was most apparent among variants lacking concordant support across the other prediction domains, for which changes in flexibility, conformational geometry, local interaction networks, or β-core organization provided additional mechanistic resolution. Low MD perturbation, in turn, more often coincided with neutral predictions across the other approaches. These patterns support dynamics-based analysis as a complementary source of structural information that can distinguish different modes of variant-associated perturbation. Importantly, prior experimental work provides a biological point of comparison. Site-directed mutagenesis identified R61, K63, K65, and K68 as determinants of hBD-1 anti-*E. coli* activity despite limited effects on overall tertiary structure [12]. Of the nine substitutions affecting these positions in the present study, seven (77.8%) exhibited at least intermediate MD perturbation, while none met the majority damaging criterion for either static predictor domain. This does not establish *S_MD_* as a predictor of antimicrobial activity but demonstrates that substitutions at experimentally implicated functional residues can exhibit measurable dynamic perturbation despite limited support from static predictors. These variants therefore emerge as prioritized candidates for direct functional testing. A focused experimental panel spanning high-*S_MD_*variants, low-*S_MD_* controls, C-terminal charge substitutions, and representatives of the major disulfide-linked perturbation mechanisms could test whether modeled signatures correspond to changes in antimicrobial function.

The sequence-distal flexibility responses observed for several variants suggest that hBD-1 perturbations can extend beyond the substitution site. Recurrent changes at residues distant in primary sequence are consistent with conformational coupling within the compact defensin fold, where local alterations may redistribute structural constraints across the peptide. Such effects may be particularly relevant for substitutions outside highly conserved motifs, for which local sequence context alone may be insufficient to predict molecular consequences.

Cysteine substitutions provide a clear example of this mechanistic heterogeneity. Although disulfide-forming cysteines were consistently recognized as important across prediction approaches, the modeled consequences of their substitution were not uniform. Individual substitutions instead produced distinct effects on flexibility, local contact organization, or β-core maintenance. These findings suggest that loss of a disulfide-forming residue can remodel hBD-1 through multiple mechanisms depending on its position and the biochemical properties of the substituted amino acid. This heterogeneity may help explain why variants affecting similarly conserved residues do not necessarily produce equivalent molecular behavior. More broadly, these findings emphasize that conservation identifies structurally important positions but does not, by itself, specify the molecular mechanism through which a particular substitution may act.

## Conclusion

The integrated sequence-, structure-, and dynamics-based analyses provide complementary information about distinct molecular properties and resolve mechanistic differences among variants that appear similar when considered through a single analytical lens. The resulting variant-specific perturbation signatures provide a basis for prioritizing functional studies without assigning variants to binary functional or dysfunctional categories. The dynamics-perturbation classes used here represent relative prioritization within the analyzed variant set rather than experimentally validated functional thresholds. The MD results are influenced by the modeled starting structure, force field, solvent representation, simulation length, and scoring definitions. The simulations also evaluate intrinsic peptide behavior and do not capture the full biological environment in which hBD-1 functions, including membrane interactions, receptor engagement, secretion, or other cellular processes. Because antimicrobial function was not measured, *S_MD_* should be interpreted as a quantitative summary of modeled perturbation and a hypothesis-generating tool, not as a measure of clinical risk or antimicrobial potency. Within these boundaries, further evaluation is needed to determine whether these modeled perturbation patterns correspond to altered antimicrobial activity or receptor engagement, and whether they are associated with genotype-phenotype relationships in clinical settings characterized by increased infection susceptibility. Together, this work establishes a molecular foundation for investigating how human hBD-1 coding variation may influence peptide behavior and host defense in biologically relevant renal, urogenital, and transplant settings.

## Supporting information

Supplemental Equations

## Supplementary Materials

The following supporting information can be downloaded with this article: Figure S1: Potential-energy trajectories for WT hBD-1 and representative variants; Figure S2: Multiple sequence alignment of human β-defensins; Figure S3: Variant-specific RMSF perturbations and identification of flexibility hotspots in hBD-1; Figure S4: MD-derived features of the hBD-1 variant cohort; Equations S1-S14: Detailed definitions of MD-derived scoring metrics.

## Author Contributions

Conceptualization, R.U., J.C.H., and G.L.; Methodology, R.U. and R.J.F.; Formal Analysis, R.U. and R.J.F.; Data Curation, R.U., F.P.V., and R.J.F.; Writing - Original Draft Preparation, R.U., R.J.F., and G.L.; Writing - Review & Editing, F.P.V., J.C.H., R.U., R.J.F., and G.L.; Visualization, R.J.F.; Funding Acquisition, J.C.H, R.U., and G.L.

## Institutional Review Board Statement

The study was conducted in accordance with the Declaration of Helsinki and approved by the Institutional Review Board of Mayo Clinic (IRB 19000001).

## Informed Consent Statement

Informed consent was obtained from all subjects involved in the study.

## Data Availability Statement

Additional data may be made available upon request to the authors.

## Acknowledgments

The authors thank the Mayo Clinic Tapestry team and patients. During the preparation of this manuscript, the authors used OpenAI ChatGPT as a support tool for developing and troubleshooting data-analysis scripts and for limited manuscript editing. All analyses were carried out by the authors and executed independently. The authors have reviewed and edited the output and take full responsibility for the content of this publication.

