## Supplemental Equations for "Structural and Molecular Dynamics Analysis of Human β-Defensin 1 Variants Reveals Distinct Perturbation Signatures"

### **Supplementary Material: Equations Used for Score Derivation**

**Notation.** $v$, variant; WT, wild type; $r$, residue; $R$, number of analyzed residues ($R=45$); $j$, independent simulation replicate; $i$, contact; $F$, analyzed frames ($F=25$); $B$, selected $\beta$-core residues ($B=10$); $\bar{x}$, mean across the 10 replicate-level values; $s_{\mathrm{WT}}$, sample standard deviation across the 10 WT replicate-level values; $I\left( \cdot\right)$, indicator function.

#### Generic WT-standardization and bounded component score

$$\Delta X_{v}=\bar{X}_{v}-\bar{X}_{\mathrm{WT}} \left( S1 \right)$$

$$Z_{X,v}^{\mathrm{WT}}=\frac{\Delta X_{v}}{s_{\mathrm{WT}}\left( X \right)} \left( S2 \right)$$

$$S_{X,v}=min\left( \frac{\left| Z_{X,v}^{\mathrm{WT}} \right|}{2},1 \right) \left( S3 \right)$$

Here, $Z_{X,v}^{\mathrm{WT}}$ denotes the WT-standardized difference. An absolute difference of two or more WT replicate standard deviations maps to a component score of 1.

#### RMSF perturbation score

$$\Delta\mathrm{RMSF}_{v,r}^{\mathrm{WT}\text{-}\mathrm{std}}=\frac{{\overline{\mathrm{RMSF}}}_{v,r}-{\overline{\mathrm{RMSF}}}_{WT,r}}{s_{WT,r}\left( \mathrm{RMSF} \right)} \left( S4 \right)$$

$$H_{v,r}=I\left( \left| \Delta\mathrm{RMSF}_{v,r}^{\mathrm{WT}\text{-}\mathrm{std}} \right|\geq2 \right) \left( S5 \right)$$

$$S_{\mathrm{RMSF}\left( \mathrm{WT}\text{-}\mathrm{std} \right),v}=min\left[ \frac{\frac{1}{R}\sum_{r=1}^{R} \left| \Delta\mathrm{RMSF}_{v,r}^{\mathrm{WT}\text{-}\mathrm{std}} \right|}{2},1 \right] \left( S6 \right)$$

$$S_{\mathrm{RMSF}\left( \mathrm{breadth} \right),v}=min\left[ \frac{\frac{1}{R}\sum_{r=1}^{R} H_{v,r}}{0.25},1 \right] \left( S7 \right)$$

$$S_{RMSF,v}=\frac{S_{\mathrm{RMSF}\left( \mathrm{WT}\text{-}\mathrm{std} \right),v}+S_{\mathrm{RMSF}\left( \mathrm{breadth} \right),v}}{2} (S8)$$

#### Conformational-remodeling score

$$S_{Rg/SASA,v}=\frac{S_{Rg,v}+S_{SASA,v}}{2} \left( S9 \right)$$

$S_{\mathrm{Rg}}$ and $S_{\mathrm{SASA}}$ are each calculated using Equations S1–S3 after averaging frames within each independent replicate.

#### Contact-remodeling score

$$J_{w,v}=\frac{\sum_{i} \min\left( w_{WT,i},w_{v,i} \right)}{\sum_{i} \max\left( w_{WT,i},w_{v,i} \right)} \left( S10 \right)$$

$$S_{contact,v}=1-J_{w,v} \left( S11 \right)$$

$w$ denotes mean contact occupancy; contacts absent from one profile are assigned an occupancy of 0.

#### Secondary-structure and fold-integrity scores

$$O_{v,j}^{E}=\frac{N_{E,v,j}}{B\times F} \left( S12 \right)$$

$N_{E,v,j}$ is the number of DSSP E-state assignments across the $B$ selected $\beta$-core residues and $F$ analyzed frames for replicate $j$.

$$S_{ss,v}=min\left[ \frac{\left| \left( \bar{O}_{v}^{E}-\bar{O}_{\mathrm{WT}}^{E} \right)/s_{\mathrm{WT}}\left( O^{E} \right) \right|}{2},1 \right] \left( S13 \right)$$

$$S_{fold,v}=\frac{S_{contact,v}+S_{ss,v}}{2} \left( S14 \right)$$
