## Supplementary figures and images for "Structural and Molecular Dynamics Analysis of Human β-Defensin 1 Variants Reveals Distinct Perturbation Signatures"

### Figure S1

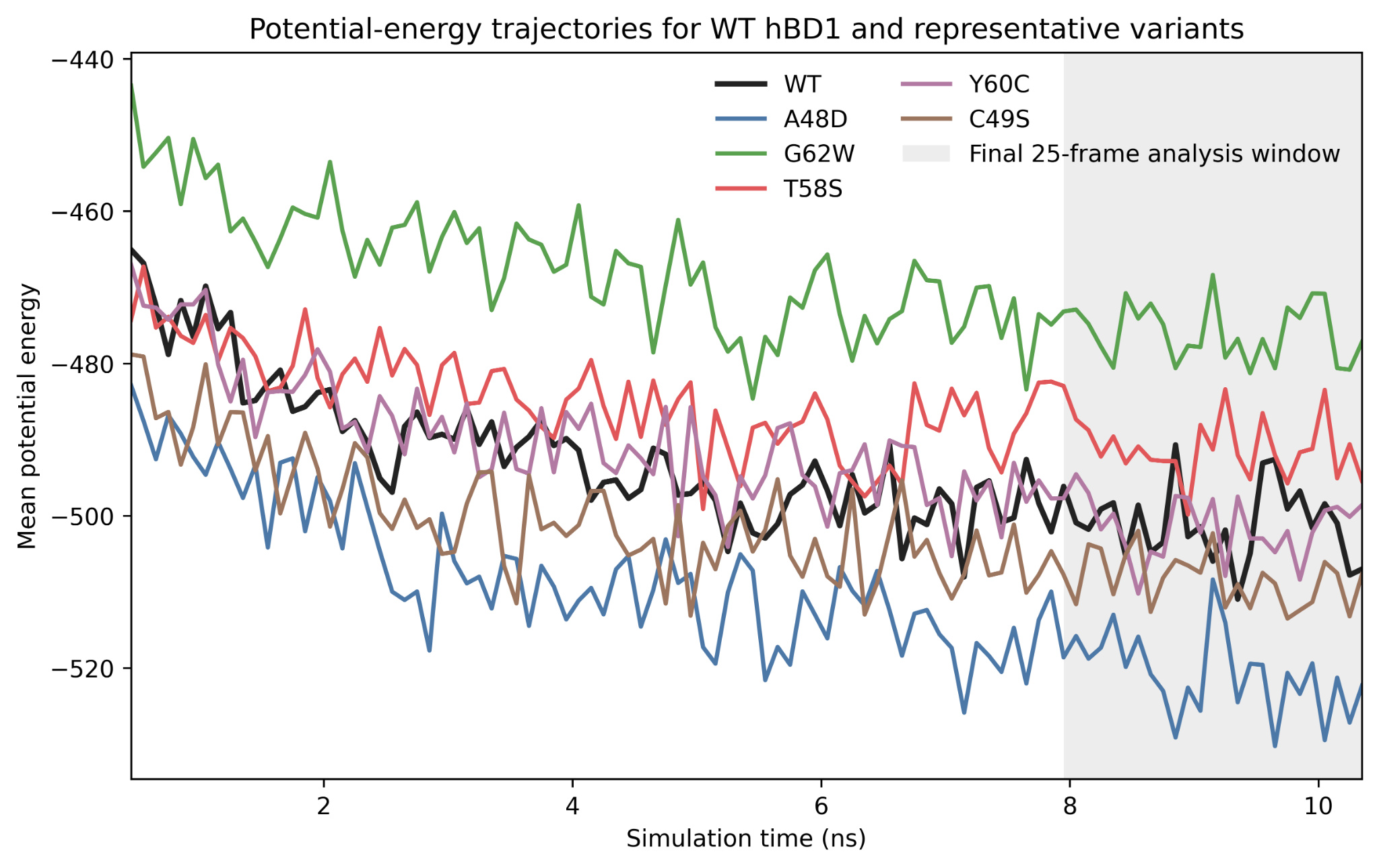

### Figure S2

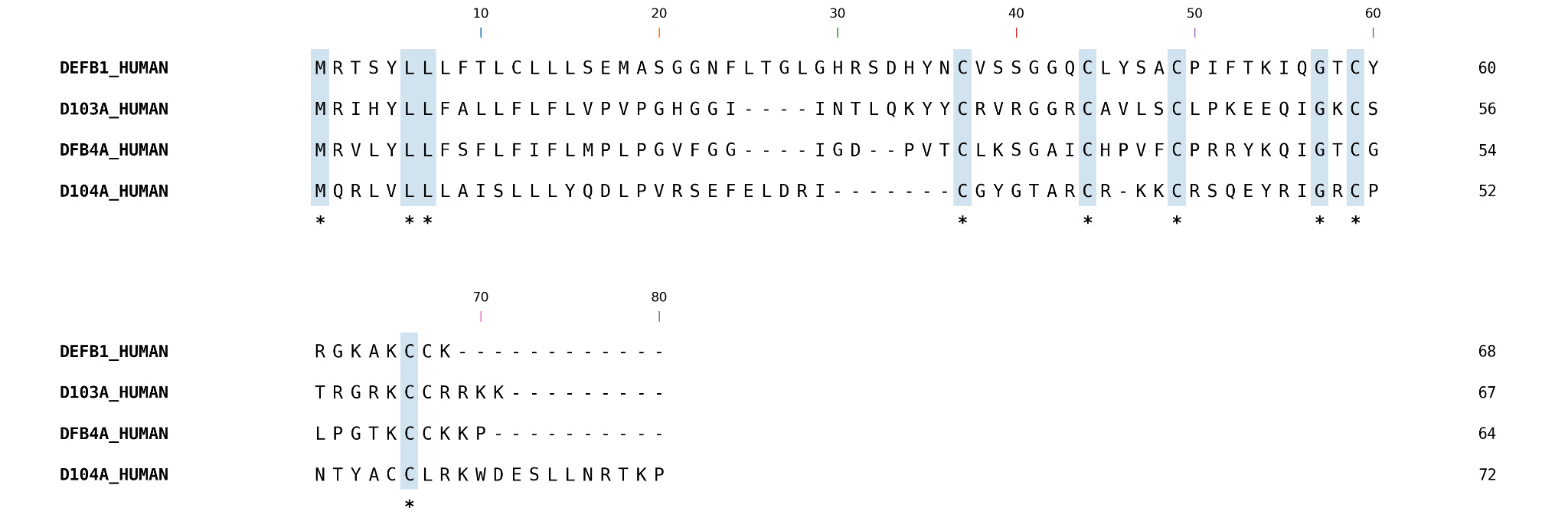

### Figure S3

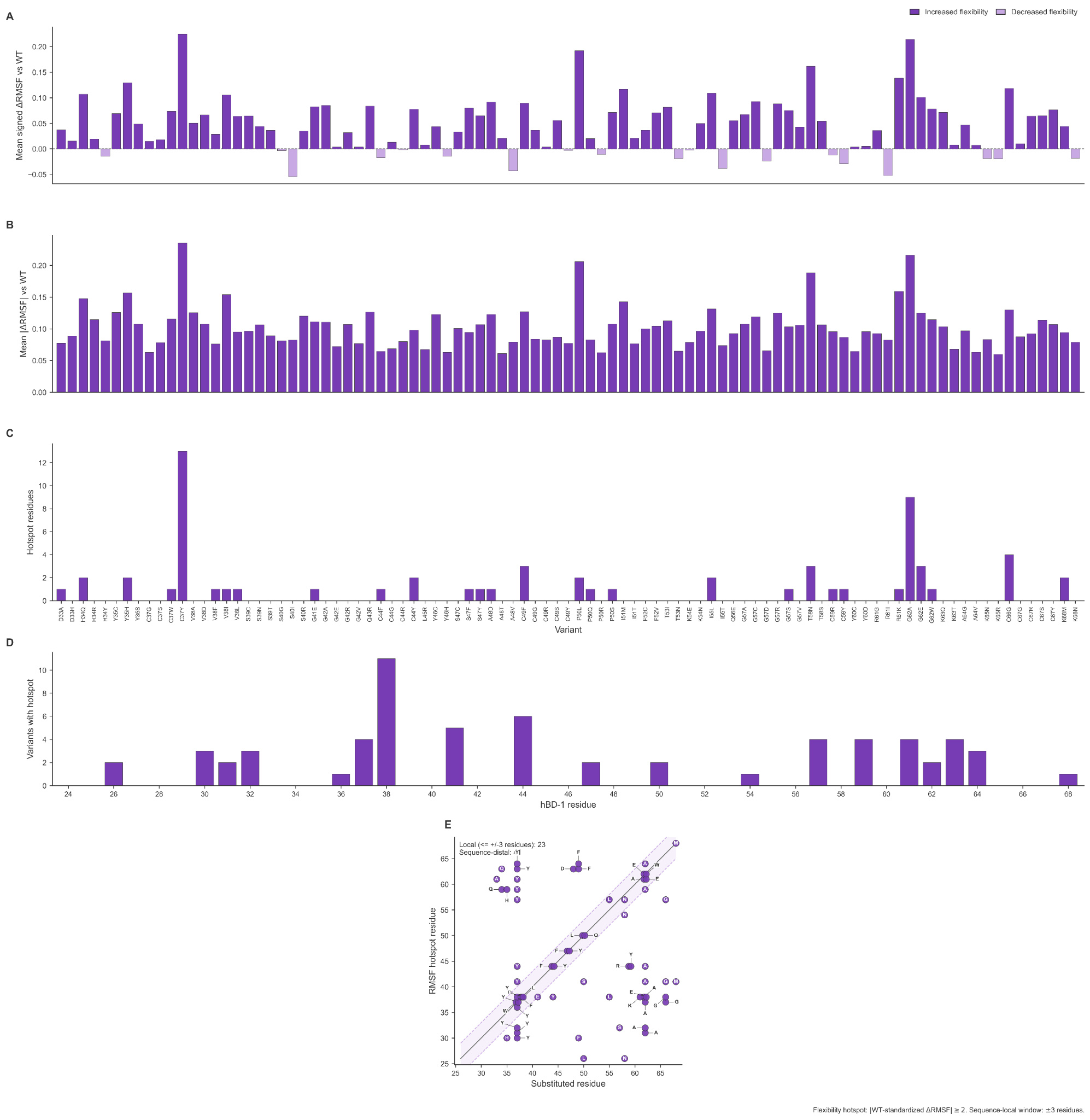

### Figure S4

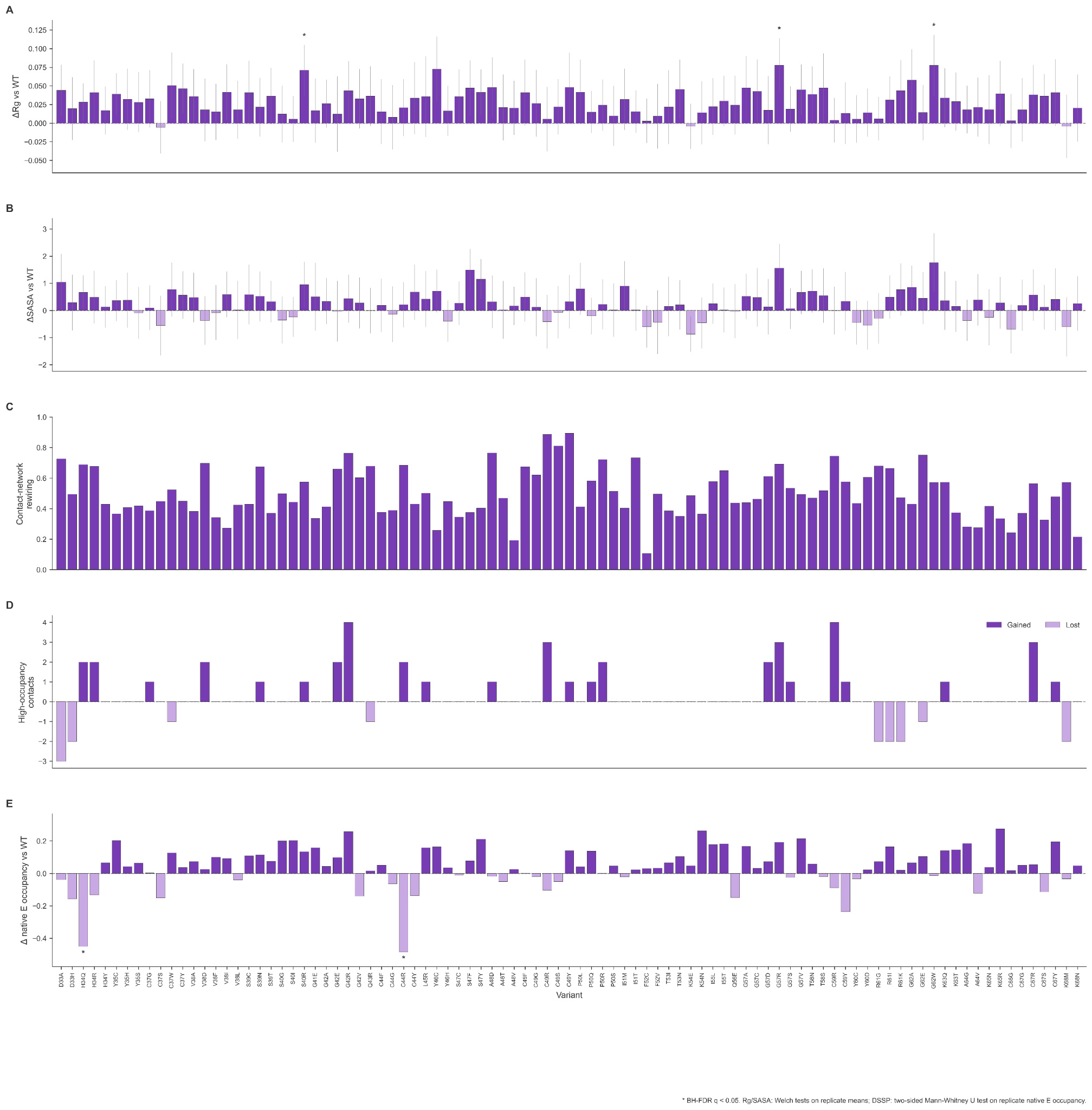
